# Relationship between Protein Conformational Stability and Its Immunogenicity When Administering Antigens to Mice Using Adjuvants

**DOI:** 10.1101/2023.10.26.564131

**Authors:** Kosuke Oyama, Tadashi Ueda

## Abstract

Antigen-presenting cells (APCs) are crucial in the immune system by breaking down antigens into peptide fragments that bind to major histocompatibility complex molecules. Previous research suggests that stable proteins may hinder CD4^+^ T cell stimulation by impeding antigen processing and presentation. Conversely, some proteins require stabilization to activate the immune response. This discrepancy may be influenced by various factors, including protein characteristics and the use of different adjuvants in animal experiments. Here, we investigated the effects of adjuvants on antigen administration, specifically focusing on the stability of the CH_2_ domain. Together with the previous study, we show that protein stability is also crucial in triggering an immune response in mice by binding protein antigens to B cell receptors on APCs. Together with the study so far, we propose that intrinsic protein stability is crucial for binding to B cell receptors on APCs in mice when administering antigens with adjuvants.

## Introduction

Vaccines are created to stimulate the immune system. In vaccine development, vaccine immunogenicity can be enhanced through modifications of protein antigens, such as amino acid mutation [1] or chemical modification [2], as well as oligomerization of protein antigens [3]. Protein digestion by proteases is faster when protein conformational stability is lower [4–7]. This highlights the close relationship between protein conformational stability and vaccine development [8,9]. Antigen-presenting cells (APCs) utilize proteases to degrade antigens into peptide fragments, which subsequently bind to major histocompatibility complex (MHC) molecules. The peptide-MHC complexes are presented on the cell surface and recognized by T cells. This interaction activates T cells and initiates an immune response against the specific antigen, thereby triggering immune responses [10–13]. There is evidence suggesting that stable proteins have a lower ability to stimulate CD4 positive T cells, such as helper T cells [14–17], and/or antibody responses [14,17–20], because of inefficient antigen processing and presentation. On the other hand, some proteins can only trigger an immune response when they are stabilized [21,22]. Therefore, the strategy to enhance the immune response to protein antigens by manipulating protein conformational stability in vaccine design is still not fully understood [8,9]. These experiments employed different proteins such as charges, molecular weight, intrinsic conformational stability and different adjuvants in the administration of antigens into mice. We have recently shown that mice immunized with the SARS-CoV-2 spike protein mixed with Alum produced neutralizing antibodies in sera. Sera from mice immunized with the protein mixed with CFA/IFA did not produce an effective neutralizing response, even though both adjuvants inducing an IgG response [23]. Under the circumstances, in this study, our focus is on the various adjuvants that have been previously used to administer antigens to mice.

IgG is a dimeric glycoprotein displaying a conserved N-glycosylation site in each of its CH_2_ domains in the effector region, referred to as Fc [24]. The CH_2_ domain in human IgG1 is typically unstable [25], but its conformational stability can be increased by 20°C through the introduction of a disulfide bond using site-directed mutagenesis using *E.coli* expression and *in vitro* refolding, in which the thermally increased CH_2_ domain is referred to as mut20 [26]. Therefore, the CH_2_ domain was anticipated to be a promising scaffold for developing new therapeutics. We successfully produced the CH_2_ domain from *Pichia pastoris* at a high yield [27]. Additionally, we showed that the retardation of aggregation of the CH_2_ domain by combining glycosylation and its stabilization by introducing a disulfide bond [28], in which the disulfide bond used in this study is the same as the one reported in a previous study [26]. As protein aggregations are reported to be closely related to immunogenicity [29,30], in this study, we investigated the immune responses to the administration of the human CH_2_ domain, which has moderate stability (melting temperature is approximately 50°C [28], and its stabilized variant (mut 20) derived from *Pichia pastoris* in the presence of two types of adjuvants, CFA/IFA and Alum. As a result, we happened to find the differences in immune responses when administering the human CH_2_ domain and its stabilized variant in the presence of two types of adjuvants, CFA/IFA and Alum. Namely, the CH_2_ domain elicited a stronger IgG response compared to its stabilized counterpart when PBS and Alum were used. On the other hand, animal experimentation using CFA/IFA yielded contrasting results. In addition to studying immune responses [16–21], we also investigated the correlation between protein conformational stability and immunogenicity when administering antigens to mice using adjuvants in order to design antigens for vaccine development.

## Materials and Methods

### Materials and Reagents

The CH_2_ domain and mut20, which are the human CH_2_ domain and mut20 without glycosylation from *Pichia pastoris*, were prepared according to our previous study [27,28]. Hen lysozyme and ribonuclease A were purchased from Nacalai Tesque, Inc. (Japan). Cathepsin B and ribonuclease S were purchased from Sigma Aldrich (USA). Reduced and carboxymethylated lysozyme was prepared according to the previous study [31]. All other chemicals employed in the study were of the highest quality commercially available.

### Mice

Thirty female BALB/c mice were obtained from Japan SLC, Inc. (Japan) and immunized at seven weeks of age. All animal experiments were conducted according to the relevant national and international guidelines in the Act on Welfare and Management of Animals (Ministry of Environment of Japan) and the Regulation of Laboratory Animals (Kyushu University) and under the protocols approved by the Institutional Animal Care and Use Committee review panels at Kyushu University.

### Immunizations

Five mice were administered the CH_2_ domain or mut20 (30 μg/shot) dissolved in Alum adjuvant or CFA adjuvant intraperitoneally (i.p.) or the protein samples (100 μg/shot) dissolved in PBS at 0 days. On days 7 and 14, the mice received i.p. booster immunizations with the same protein (30 μg/shot) dissolved in Alum adjuvant or IFA, or with the same protein (100 μg/shot) dissolved in PBS. Blood samples were drawn every week. Serum antibodies were detected using ELISA.

### ELISA

96-well ELISA plates were coated with CH_2_ domain or mut20 in coating buffer (1.59 g sodium carbonate, 2.93 g sodium bicarbonate, 0.2 g sodium azide, 1 L of distilled water, pH 9.6), at a concentration of 1 µg/ml and a temperature of 4°C overnight. After washing with PBST, blocking was achieved with a solution containing 5% skim milk in PBST at 37°C for one hour. After washing with PBST, 100 µl of mouse sera at various concentrations were incubated in the plates at 37°C for one hour. After washing with PBST, plates were incubated with 1:10000-diluted horseradish peroxidase-conjugated goat Fab anti-mouse IgG solution at 37°C for one hour. After washing with PBST, 50 µl of ABTS solution (0.3 mg 2,2’-azino-bis (3-ethylbenzothiazoline-6-sulfonic acid) diammonium salt in 20 ml of 0.1 M citrate buffer containing 0.04 µl of 30% H_2_O_2_, pH 4.0) was added to the plates. The plates were incubated at 37°C and the absorbance was measured at 405 nm.

### Protease digestion assay

Fifty micrograms of protein substrates were prepared to 0.5 mg/ml in 100 mM MES buffer (pH 5.0) and 5 mM mercaptoethanol. The substrates were added to 0.2 mU cathepsin B, a lysosomal cysteine protease. Digestion reactions were performed at 37°C and stopped at various time points by freezing the samples. After centrifugation (13,000 rpm, 4°C, 10 min), the analysis of the digested peptide in the supernatant was performed by size exclusion chromatography (SEC) using a column of PolyHYDROXYETHYL A column (200 Å, 5 μm, 4.6 mm × 200 mm, PolyLc inc., USA) at room temperature and a flow rate of 0.1 ml/min. The mobile phase buffer was composed of 200 mM Na_2_SO_4_ and 5 mM KH_2_PO_4_ supplemented with 25% acetonitrile (pH 3.0)

### MALDI-TOF-Mass

Protease digests were prepared according to the protease digestion assay. Digestion reactions were performed at 37°C for 24 hr. After cathepsin digestion, the samples were desalted using Zip tip C18. Desalted samples were eluted directly to a target plate and air-dried. The samples were covered with 1 μl α-cyano-4-hydroxycinnamic acid saturated solution in 50% acetonitrile and 0.1% trifluoroacetic acid (TFA) and air-dried. MALDI-TOF-Mass was performed using an AutoflexⅢ mass spectrometer (Bruker Daltonics, Germany) with sinapinic acid as a matrix reagent.

## Results and Discussion

### The impact of adjuvants on the immunogenicity of the CH_2_ domain with varying conformational stabilities

In order to investigate the relationship between the stability and immunogenicity of the CH_2_ domain, two types of adjuvants were used. The CH_2_ domain or mut 20 in PBS (without adjuvant) were administered to mice, along with Alum or CFA/IFA, following the protocol (Fig. 1A). The presence of anti-CH_2_ domain or anti-mut20 antibodies in the serum of immunized mice was detected using ELISA. In animal experiments, the immune response (IgG production) elicited by mut20 in mice was lower than that of the intact CH_2_ domain when Alum was used as an adjuvant or when no adjuvant (PBS) was used (Fig. 1B). Unexpectedly, when CFA/IFA was used as an adjuvant, the immune response was contrary to what was expected. The production of IgG in mice induced by mut20 was higher than that induced by the intact CH_2_ domain (Fig. 1B).

**Fig. 1.**
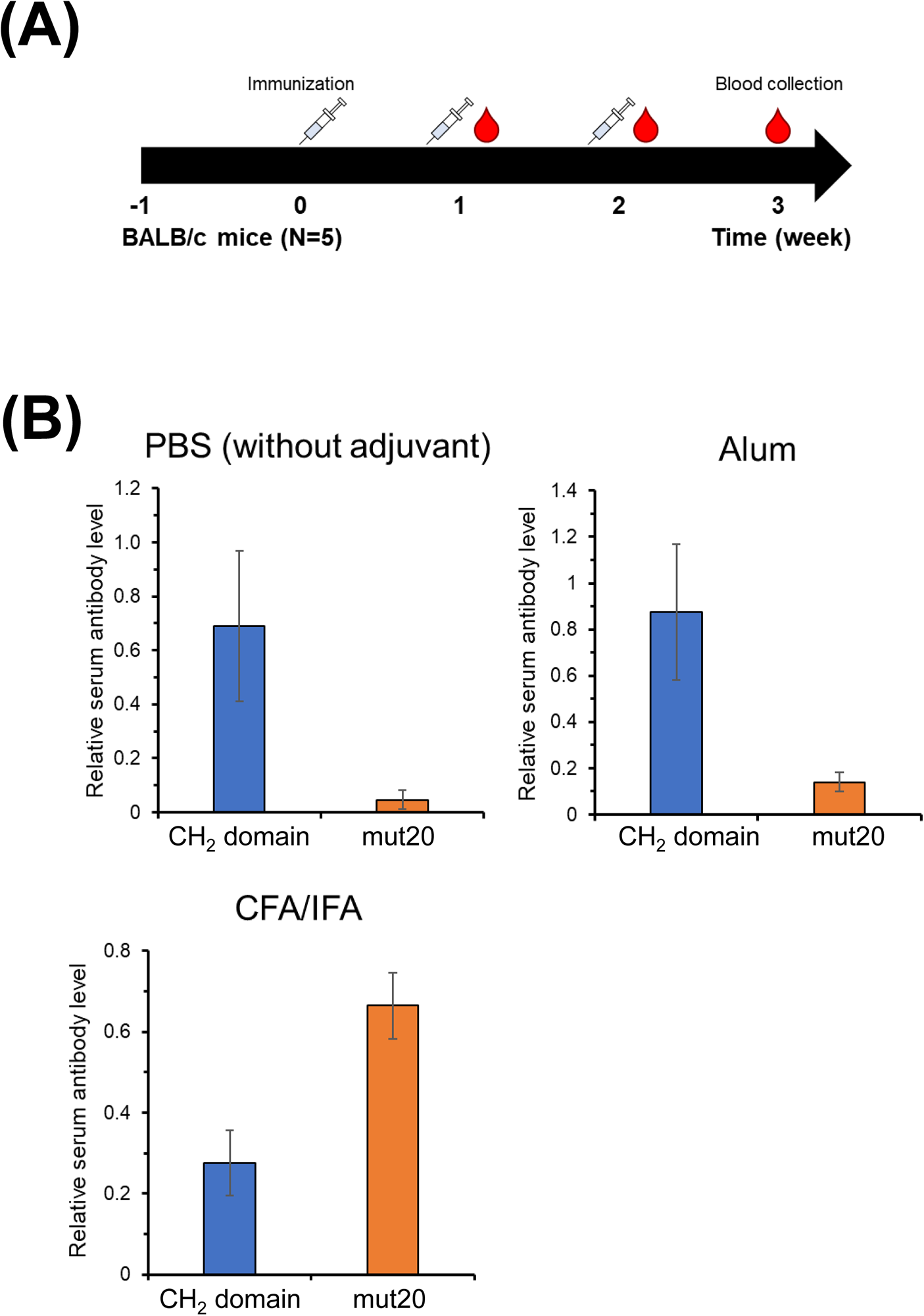
(A) Schematic representation of the immunization schedule for BALB/c mice. CH_2_ domain and mut20 were inoculated into mice with either CFA/IFA or Alum adjuvant (30 μg/shot) or PBS (100 μg/shot). Mice were boosted at 1 and 2 weeks, respectively. Serum samples were collected at 1, 2, and 3 weeks after the initial vaccination. (B) ELISA analysis was performed to detect the increase of IgGs bound to the CH_2_ domain or mut20 in the sera of vaccinated mice 3 weeks after the first immunization (N = 5). The CH_2_ domain or mut20 was coated onto ELISA plates.

### Protease digestion assay of CH_2_ domains

Immune responses have been found to be closely linked to the resistance of proteases in APCs [14–17]. The analysis focused on the amount and length of peptides generated by protease digestion of the CH_2_ domains. This is because the extent of the immune response is influenced by the amount of the complex formed between the MHC class II receptor and peptide on helper T cells. To assess antigen processing by APCs *in vitro*, researchers often use cathepsin B, a major protease found in lysosomes that has both endopeptidase and exopeptidase activities [32–34]. Cathepsin B is often used in studies on protein conformational stability and the immunogenicity of proteins. The antigens were digested at 37°C under reducing and acidic conditions (pH 5.0) to simulate the lysosomal environment in cells. The peptides resulting from the digestion of the CH_2_ domains were analyzed using MALDI-TOF-Mass, as stated in a previous study [32]. The MS chromatogram of the intact CH_2_ domain after cathepsin B digestion showed a similar profile to that of mut20 after cathepsin B digestion. This suggests that there was no difference in the proteolytic fingerprint between the two CH_2_ domains, except for the mutation sites in mut20 (Fig. 2A and 2B). To analyze the digestion time course, the CH_2_ domains were subjected to SEC after cathepsin digestion at various time points using a PolyHYDROXYETHYL A column. The column was able to separate tryptic peptides of reduced and S-carboxymethylated hen lysozyme. The molecular weights of 940 and 1750 corresponded to peptides of seven and sixteen lengths, respectively (Fig. 3A). Since the crystallographic analysis has shown that MHC class II on T cells can bind at least octamer peptides in their presentation to T cell receptors [35], the column could potentially be used to analyze the quantities of peptides from the CH_2_ domain or mut20 through cathepsin B digestion. These peptides would then be able to bind to the MHC class II receptor on T cells. It was found that the intact CH_2_ domain was digested by cathepsin B at a significantly faster rate than mut20, as shown in Fig. 3B and 3C. This suggests that mut20 is more resistant to the lysosomal protease action of cathepsin B. It has been controversial to determine whether the conformational protein stability of an antigen affects its immunogenicity or not [8,15–22]. Therefore, we conducted animal experiments to investigate the impact of various adjuvants on the immunogenicity of both the intact CH_2_ domain and its stabilized form (mut 20). As demonstrated in animal experiments, immunization of intact CH_2_ domain mixed with Alum or just PBS into mice resulted in a stronger IgG response compared to the stabilized form. In Fig. 3B and 3C, it is evident that the intact CH_2_ domain was more susceptible to cathepsin B digestion compared to mut20. We have confirmed that stabilizing the CH_2_ domain reduces immunogenicity by inhibiting protease digestion by cathepsin B. It was demonstrated that the immune response in mice is diminished when the conformational protein stability increases in the case of the CH_2_ domain prepared in PBS, which is commonly used in the administration of biotherapeutics. When considering the instability of the CH_2_ domain compared to other domains in human antibodies, this finding may contribute to reducing the immunogenicity of therapeutic antibodies, because protein aggregations often occur due to conformational distortion starting from the most unstable region of a protein, and this is closely related to increased immunogenicity of therapeutic antibodies [36–39]. On the other hand, when using CFA/IFA adjuvant, the intact CH_2_ domain in mice showed a lower immune response (IgG production) compared to mut20. In this case, the immune response cannot be solely explained by the amount of peptide digested by cathepsin B. It has been reported that Freund’s adjuvant can decrease the conformational stability of antigens because of its hydrophobic nature [40,41]. Therefore, in order to generate an immune response in moderately stable proteins such as the CH_2_ domain (with a melting temperature of 52.6°C at pH 5.5 [28]), additional protein stabilization is necessary to counteract destabilization caused by interactions with Freund’s adjuvant. This strategy of enhancing immunogenicity is consistent with previous studies that have shown the significance of stabilizing protein antigens such as RNase S and apo-Horseradish Peroxidase (apo-HRP), which has a lower melting temperature (Tm) than the CH_2_ domain [21,22], are necessary to elicit an immune response, even when using Alum adjuvant in animal experiments with mice.

**Fig. 2.**
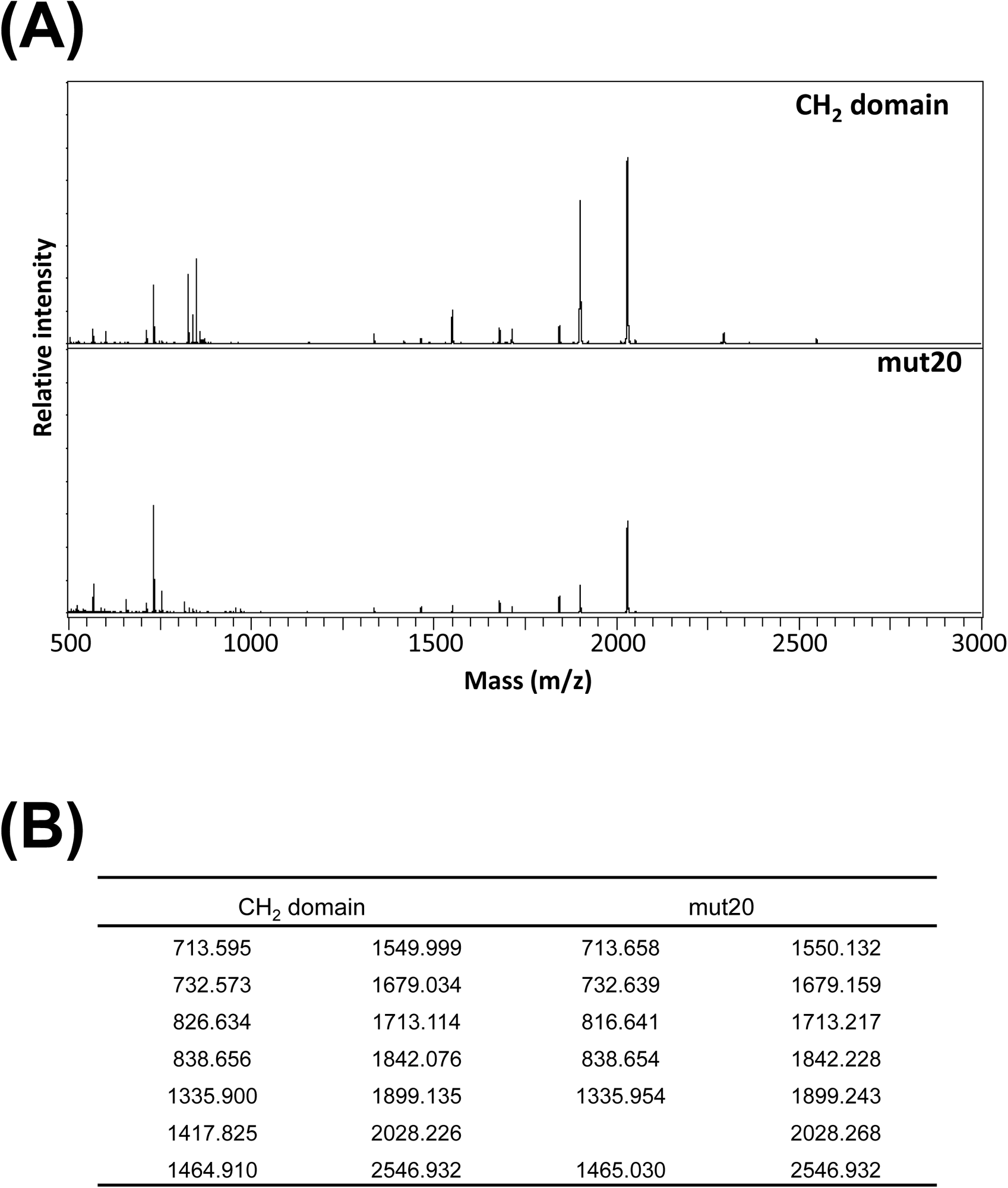
(A) MS chromatograms of the CH_2_ domain and mut20. The digestion reaction with cathepsin B was conducted at 37°C for 24 hours. The peptides were analyzed using MALDI-TOF-Mass (B) The mass of peptides is depicted in Figure 2A. The data obtained from MALDI-TOF-Mass. The masses of the CH_2_ domain and mut20 were essentially identical. The discrepancy of 9.993 between 826.634 and 816.641 could be attributed to the substitution of leucine and cysteine as predicted by the peptide mass analysis.

**Fig. 3.**
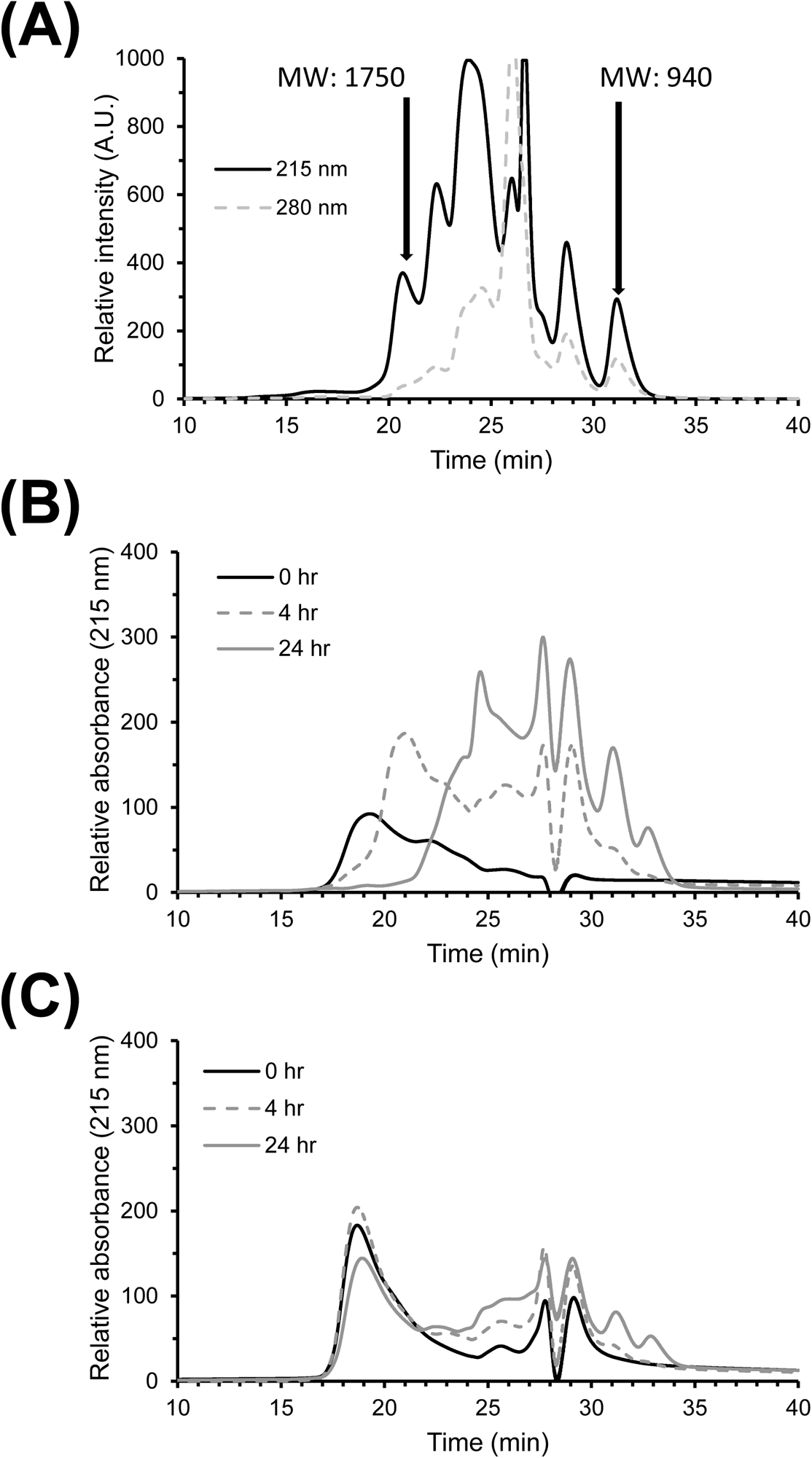
Size exclusion chromatography (SEC) using a column of PolyHYDROXYETHYL column at room temperature and a flow rate of 0.1 ml/min. (A) SEC patterns of the tryptic peptides of reduced and S-carboxymethylated hen lysozyme were monitored using a UV detector at 215 nm and 280 nm. The elution times of the tryptic peptides with molecular weights of 1750 and 940 are indicated. (B) SEC patterns of the digested peptides from the CH_2_ domain were monitored at pH 5.0 and 37°C using cathepsin B for 0 hr (black solid line), 4 hr (gray broken line) and 24 hr (gray solid line). The monitoring was done using a UV at 215 nm. (C) SEC patterns of the digested peptides of mut20 were monitored at pH 5.0 and 37°C by cathepsin B for 0 hr (black solid line), 4 hr (gray broken line) and 24 hr (gray solid line) using a UV at 215 nm.

### Protease digestion assay of ribonucleases

The Tm of RNase S without S-peptide was reported to be 38.6°C in a 10 mM Mops buffer with 200 mM NaCl at pH 7.0 [42], while Tm of RNase A was found to be 62.6°C in a 10 mM Mops buffer at pH 7 [43]. At pH 5, the Tm of RNase S without S-peptide was 43.4°C, while that of RNase A was 60.2°C [44]. The recombinant HRP (holo form) had a Tm of 68.9°C in Bis-Tris buffer at pH 7 [45], which is similar to the Tm of intact HRP (holo form) at 70°C in 100 mM phosphate buffer at pH 7.0 [46]. In Bis-Tris buffer at pH 7, the recombinant HRP (apo form) could not be evaluated because the protein unfolded before reaching the temperature of 45°C. Under the same measuring conditions (in Bis-Tris buffer at pH 7), the Tms of plant HRP in its holo form and apo form were 77.7°C and 37.2°C, respectively [45]. The order of thermal stability is as follows: plant HRP (holo form) > recombinant HRP (holo form) > plant HRP (apo form) > recombinant HRP (apo form). Thus, the Tm of recombinant HRP (apo form) is below 37.2°C, while the Tm of recombinant HRP (apo form) is 68.9°C at pH 7. A similar trend of significant instability was observed in the other HRP when heme was removed, as shown in a study that found the apo form of soybean peroxidase (SBP) has a Tm of 38°C, while the holo form of SBP has a Tm of 75°C at pH 7.0 [47].

Since previous reports indicated that the Tm of RNase S was higher than that of apo-HRP, our focus is on the protease digestion of RNase S using cathepsin B, with RNase A as a control under similar conditions as in the present study. As shown in Fig. 4, RNase A showed higher resistance to protease digestion compared to RNase S due to its higher conformational stability at pH 7 [42,43] and pH 5 [44], as previously mentioned. The amount of digestion peptides from RNase S was higher than that from RNase A, as depicted in Fig. 4B and C. It has been previously reported that RNase S is more susceptible to protease digestion compared to RNase A, moreover, RNase A is more immunogenic than RNase S [21,22]. The peptide-MHC complexes are presented on the cell surface and recognized by T cells. This interaction activates T cells and initiates an immune response against the specific antigen, thereby triggering immune responses [10–13]. When examining the MHC-peptide complex in APCs and T cell presentation, it is suggested that the immunogenicity of RNase A compared to RNase S is not solely determined by the degree of protease digestion of the RNases.

**Fig. 4.**
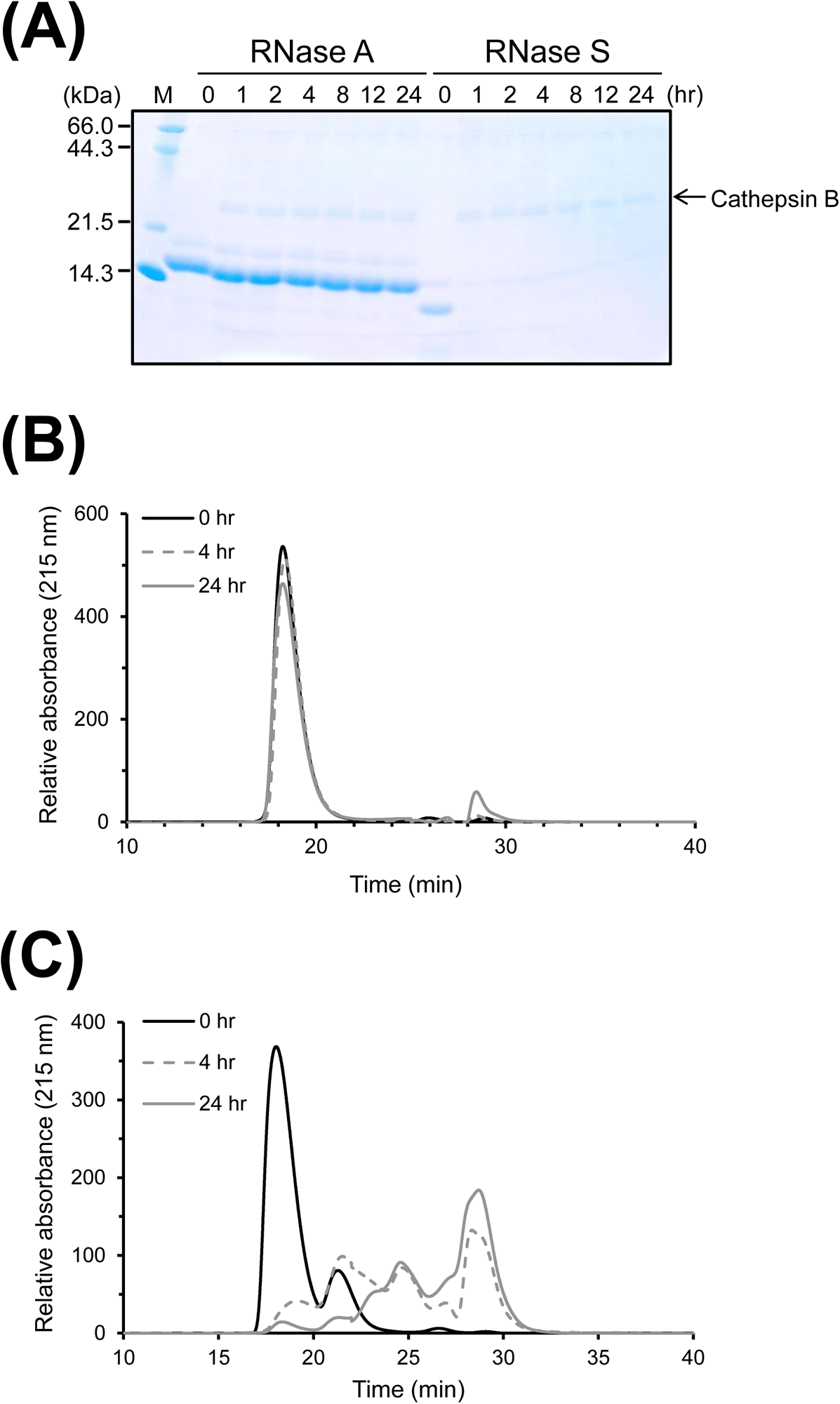
(A) SDS-PAGE results of RNase A and RNase S after cathepsin B digestion. The levels of RNase A and RNase S were analyzed after incubation with cathepsin B at pH 5.0 and 37°C for a suitable duration. SDS-PAGE followed by coomassie brilliant blue staining was used for analysis. (B) Size exclusion chromatography (SEC) patterns of the digested peptides of RNase A at pH 5.0 and 37°C by cathepsin B were monitored using a UV at 215 nm. The patterns were observed at 0 hr (black solid line), 4 hr (gray broken line) and 24 hr (gray solid line). (C) SEC patterns of the digested peptides of RNase S were monitored at pH 5.0 and 37°C by cathepsin B for 0 hr (black solid line), 4 hr (gray broken line) and 24 hr (gray solid line) using a UV at 215 nm. The chromatographic conditions for SEC were the same as shown in Figure 3.

### The intrinsic conformational stability of a protein antigen is another factor that affects its immunogenicity

In this study, we found that the immune response elicited by mut20, which is higher than at 20°C in the intact CH_2_ domain, was lower than that of the intact CH_2_ domain with a Tm of 52.5°C [28] when PBS (without adjuvant) and Alum were used as adjuvants. However, the immune response was the opposite when CFA/IFA was used as an adjuvant (Fig. 1B). Additionally, when subjected to protease digestion by cathepsin B, the CH_2_ domain was cleaved faster than mut 20 was (Fig. 3B and 3C), resulting that the difference in immune responses in Fig. 2B when using PBS (without adjuvant), Alum, and CFA/IFA could not be explained solely by the MHC-peptide complex view of APCs to T cell presentation.

The relationship between the conformational stability of a protein antigen and its immunogenicity has been studied. It was found that the immune responses in mice were reduced when protein antigens with higher melting temperatures (such as hen lysozyme, Phlp 7, Der p2 and toxin α) had increased conformational stabilities, even when using CFA/IFA or Alum as adjuvants [16–21] (in the case of “high stability” in Fig.5). According to reports, the Tm of hen lysozyme is 75°C (at pH 4.0, [48]), 69°C (in its concentrated solution at pH 4.0, [49]), 72°C (in its concentrated solution at pH 7.0, [49]), the Tm of Phlp 7 is 77.3°C (at pH 7.4, [50]), the Tm of Der p2 is above 70°C (at pH 5.0, [51]), and the Tm of toxin α is above 85°C (at pH 7.0, [16]) and above 80°C (at pH 4.3, [16]). On the other hand, the Tm of RNase S is 38.6°C (in 10 mM MOPS buffer containing 200mM NaCl at pH 7.0, [43]), and the Tm of apo-HRP is below 37.2°C (at pH 7, [45]), indicating that these proteins were intrinsically less stable than the CH_2_ domain employed in this study. As a result, it was reported that the immune responses in the administration of protein antigens with lower melting temperature (such as RNase S and apo-HPR) to mice increased when their conformational stabilities increased even when using Alum as an adjuvant [21,22].

**Fig. 5.**
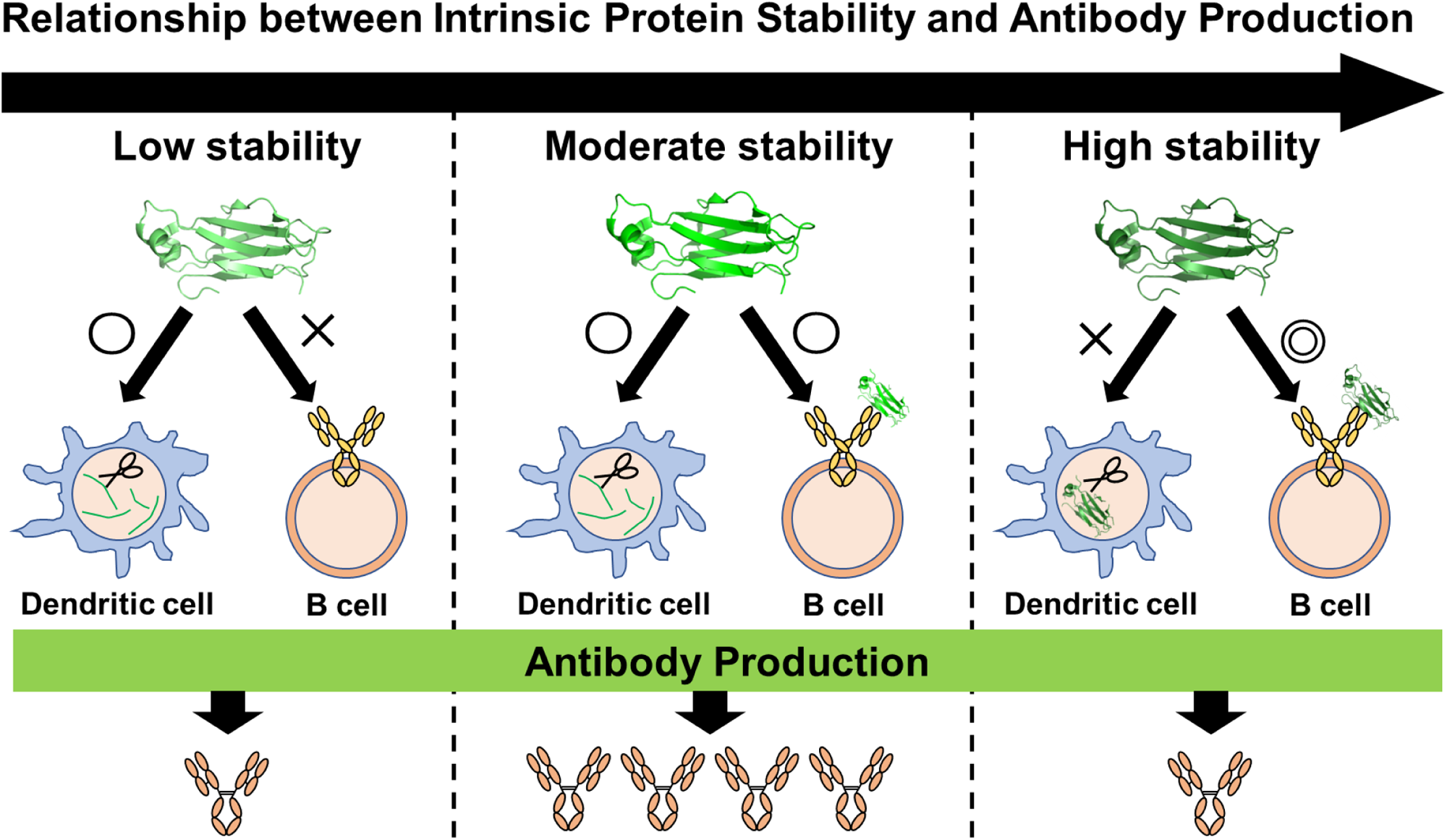
New interpretations on the relationship between protein conformational stability and antibody production.

Both the stability of protein antigens and B cell receptors (BCRs) are crucial for the binding interactions that initiate immune responses. When a BCR encounters an antigen that has a compatible conformation, the antigen binds to a specific site on the BCR, similar to a lock and key mechanism. Based on our findings, we have demonstrated that a protein antigen must possess a minimum level of intrinsic conformational stability in order to effectively bind with BCRs and induce an immune response (in the case of “low stability” in Fig.5). Indeed, the tertiary structure of RNase S starts to unfold at 37°C and completes around 52°C (with a Tm of 46°C) at pH 6.8, as observed by near-UV CD [47]. Therefore, when the population of tertiary structure in RNase S was lower in mice, it was not always able to bind the corresponding BCRs effectively, leading to reduced immunogenicity. In order to stimulate greater immune responses, it is natural to stabilize the conformational stability of RNase. Similarly, the presence of Cu^2+^ has been shown to significantly enhance vaccine immunogenicity. This is because metal ions can act as “fasteners” in macromolecules, facilitating stable formation [48]. This applies to the strategy that holo-HRP, which contains heme, is more stable than apo-HRP [43], resulting in holo-HRP being immunogenic [21,22].

Additionally, the information obtained here is consistent with our recent reports that mice immunized with a combination of the SARS-CoV-2 spike protein and Alum produced neutralizing antibodies in their sera. In contrast, mice immunized with the protein mixed with CFA/IFA did not produce an effective neutralizing response, despite both adjuvants inducing an IgG response [23].

Protein conformational stability is essential in vaccine design, particularly in light of the current focus on vaccine development. In this study, we suggest that protein antigens must possess a minimum level of intrinsic conformational stability (Fig. 5). This level ensures that the antigen’s tertiary structure is present in an animal, allowing it to effectively bind to the corresponding BCRs and initiate an immune response. In order to enhance the immune response, it may be beneficial to improve the conformational stability of a protein antigen that is unstable in mice when administering antigens to mice.

## Acknowledgments

This work was supported by the MEXT (Ministry of Education, Culture, Sports, Science, and Technology) Grant, Kyushu University operating expenses, and under the “COVID-19 Drug and Vaccine Development Donation Account” Project from Sumitomo Mitsui Trust Bank. We would like to thank Dr. Takahashi and Dr. Caaveiro from Kyushu University for their kind guidance in the experimental procedure.

## Notes

### Competing Interest Statement

The authors have declared no competing interest.

